# Multilevel and sex-specific selection on competitive traits in North American red squirrels

**DOI:** 10.1101/104240

**Authors:** David N. Fisher, Stan Boutin, Ben Dantzer, Murray M. Humphries, Jeffrey E. Lane, Andrew G. McAdam

**Affiliations:** Department for Integrative Biology, University of Guelph, Guelph, Ontario N1G 2W1, Canada; Department of Biological Sciences, University of Alberta, Edmonton, Alberta T6G 2E9, Canada; Department of Psychology, University of Michigan, Ann Arbour, Michigan 48109-1043, USA; Department of Ecology and Evolutionary Biology, University of Michigan, Ann Arbour, Michigan 48109-1043, USA; Natural Resource Sciences, Macdonald Campus, McGill University, Ste-Anne-de-Bellevue, Québec H9X 3V9, Canada; Department of Biology, University of Saskatchewan, Saskatoon, Saskatchewan S7N 5E2, Canada

**Keywords:** multilevel selection, natural selection, North American red squirrel, selection coefficient, spatial scale, *Tamiasciurus hudsonicus*

## Abstract

Individuals often interact more closely with some members of the population (e.g. offspring, siblings or group members) than they do with other individuals. This structuring of interactions can lead to multilevel natural selection, where traits expressed at the group-level influence fitness alongside individual-level traits. Such multilevel selection can alter evolutionary trajectories, yet is rarely quantified in the wild, especially for species that do not interact in clearly demarcated groups. We quantified multilevel natural selection on two traits, postnatal growth rate and birth date, in a population of North American red squirrels (*Tamiasciurus hudsonicus*). The strongest level of selection was typically within-acoustic social neighbourhoods (within 130m of the nest), where growing faster and being born earlier than nearby litters was key, while selection on growth rate was also apparent both within-litters and within-study areas. Higher population densities increased the strength of selection for earlier breeding, but did not influence selection on growth rates. Females experienced especially strong selection on growth rate at the within-litter level, possibly linked to the biased bequeathal of the maternal territory to daughters. Our results demonstrate the importance of considering multilevel and sex-specific selection in wild species, including those that are territorial and sexually monomorphic.

Data archival: the data set is archived on Dryad (info XXX), with a five-year embargo from the date of publication.

## Introduction

Phenotypic selection measures the association between individuals’ traits and some aspect of their fitness. Measures of the strength and mode of selection provide insights into the function of specific traits (Arnold 1983) and allow for predictions of how these traits might evolve across subsequent generations (Robertson 1966; Price 1970; Lande 1979; Falconer 1981; Lande and Arnold 1983). More broadly, the thousands of estimates of selection in the wild provide general lessons about the way selection often works in nature (Endler 1986; Kingsolver et al. 2001; Smith and Blumstein 2008; Cox and Calsbeek 2009; Siepielski et al. 2009, 2013).

Almost all of these estimates consider selection as acting directly on an individual’s absolute trait value or value relative to the population mean. However, individuals often interact more closely with those in their immediate environment; for instance bird nestlings compete with their siblings for access to food brought by the parents (Werschkul and Jackson 1979; Royle et al. 1999). When ecological conditions cause individuals to interact more closely with some conspecifics than others, multilevel associations between traits and fitness can arise. Under these conditions, fitness is influenced not only by the trait value of the individual, but also the trait values of litters, broods or social groups (Goodnight et al. 1992). Such multilevel selection has been shown to be equivalent to kin-selection and “neighbour-modulated selection”, where individuals influence each other’s fitness (Grafen 1984; Queller 1992; Bijma et al. 2007; Bijma and Wade 2008; but see: van Veelen et al. 2012), and may or may not correlate with selection at the level of the individual (Goodnight et al. 1992). For instance, it might be beneficial for a chick to beg more loudly than its nestmates to receive more food from the parents, but louder nests may suffer higher predation rates. The evolutionary consequences of multilevel selection are potentially striking; higher-level selection in the same direction as individual-level selection can increase the rate of the evolutionary response, but higher-level selection in the opposite direction can retard, remove, or even reverse evolutionary response to selection (Bijma and Wade 2008).

Standard measures of selection represent how trait variation across individuals relates to among-individual variation in relative fitness. These can be measured as fitness-trait covariances (selection differential; Lush 1937; Falconer 1981) and partial regression coefficients (selection gradient; Lande 1979; Lande and Arnold 1983). For example, a selection gradient is given by:

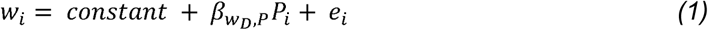

Where *w*__*i*__ is individual *i*’s relative fitness, *P*___*i*___ is *i*’s phenotype, *β*_*W*_*D*_*P*_ is the partial regression coefficient of *P*__*i*__ on *w*__*i*__, and *e*__*i*__ is a residual term. We use the notation from Bijma and Wade (2008) for consistency with later sections. The D in *β*_*W*_*D*_*P*_, indicates the effect is direct in that it is the phenotype of individual *i* influencing its own relative fitness. A single regression coefficient, *β*_*W*_*D*_*P*_, is calculated across the whole population under investigation. This implies that the component of an individual’s trait that is relevant to its relative fitness is its deviation from the population mean.

In contrast, in the context of multilevel selection, an individual’s trait can be modelled as both a deviation from its own group mean, and the deviation of the group mean phenotype from the global mean phenotype (also called “contextual analysis”; Heisler and Damuth 1987; Goodnight et al. 1992; Goodnight and Stevens 1997). An alternative is the “neighbour-modulated” or “social selection” approach, where individual phenotype values, and the mean of their neighbours (i.e. the mean of the group excluding the focal individual) are used to predict fitness (Wolf et al. 1999; McGlothlin et al. 2010). Both Queller (1992) and Bijma and Wade (2008) have shown these approaches are equivalent; we use the former for consistency with recent work on this topic by Bouwhuis et al. (2015).

Both among-individual and among-group variation may be important in determining fitness. In this case, selection is modelled with two terms: *i*’s group mean (including *i*), 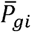, and that individual’s deviation from the group mean *ΔP*_*^i^*_ (Bijma and Wade 2008). A multilevel selection analysis can, therefore, quantify both the among-group selection gradient, 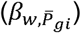, and the within-group selection gradient *β*_*W,Δ,P*_*i*__ using standard multiple regression methods for estimating selection gradients (Lande & Arnold 1983):

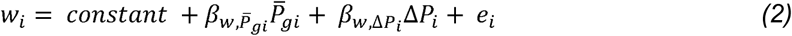

This simple two-level selection model then assumes that all groups within the population equally interact with one another. However, if some groups are clustered into a higher hierarchical level of organization (e.g. groups that share a local neighbourhood might interact more strongly) then relationships between group mean traits and group mean fitness might vary among these higher levels of organization. Therefore, the basic multilevel selection approach can be extended across any number of hierarchical levels of organization (Goodnight et al. 1992; Bijma et al. 2007).

Whilst debate over multilevel selection continues (Gardner 2015; Goodnight 2015), empirical data for its action is gathering. For example, Bouwhuis et al. (2015) found covariance between fledging mass and survival at the between-year, within-year and within-brood levels in great tits (*Parus major*), with the covariance being strongest at the broadest scales. Similarly, selection has been observed at various different levels in different systems, including among honey bee (*Apis mellifera*) colonies (Page and Fondrk 1995), among pairs of monogamous collared flycatchers (*Ficedula albicollis*) (Björklund and Gustafsson 2013), among pens of captive Japanese quail (*Coturnix japonica*) (Muir et al. 2013), among groups of jewelweed plants (*Impatiens capensis*) (Stevens et al. 1995), while contrasting individual and group-level selection was observed in water strider (*Aquarius remigis*) groups (Eldakar et al. 2009, 2010).

These examples portray organisms interacting in relatively clearly defined groups, yet animals do not always interact in such discrete units. For example populations of territorial animals consist of individuals aggregated at a range of spatial scales, from individual territories, to groups of neighbouring territories to entire populations (Coulson et al. 1997). Selection presumably could act at each of these levels simultaneously, and possibly in differing directions, but this is rarely investigated. Laiolo and Obeso (2012) found there was disruptive selection at the level of the individual for song repertoire in Dupont’s lark (*Chersophilus duponti*), but when selection on “neighbourhoods” (small populations containing 2-50 territories) was considered, selection on song repertoire was found to be stabilising. This demonstrates that non-discrete units can be a basis for selection. Nunney (1985) similarly demonstrated such “continuous arrays” of animals can be the basis for selection for altruism as they are when structured in “trait groups”.

Therefore, the key question is not whether multilevel selection is possible, but its form and strength across systems in the natural world (Biernaskie and Foster 2016). Aggregating estimates that included scales at which there might be no genetic variance in the trait might lead to an under-estimation of evolutionary change (if estimates cancel out as they are in opposing directions) or an over-estimation of evolutionary change (if the levels of selection are in the same direction). This may help us explain the inaccuracy of our predictions of evolutionary responses to selection on heritable traits (Merilä et al. 2001). Additionally, sexually antagonistic selection is quite common, and may also pose a constraint on evolution (Cox and Calsbeek 2009). However, it is unknown whether this antagonistic selection extends to multiple levels.

To study multilevel selection in an animal interacting in non-discrete groups, we focused on recruitment in a wild population of North American red squirrels (*Tamiasciurus hudsonicus*, hereafter “red squirrels”). Red squirrels defend exclusive, food-based territories centred on a cache of hoarded white spruce (*Picea glauca*) cones (Smith 1968). Most of the variation in lifetime reproductive success is determined by whether or not squirrels acquire a territory during their first year before winter commences (McAdam and Boutin 2003b; McAdam et al. 2007). Juveniles cannot oust adults from their territories, so they must find vacant territories or, if resource availability is high, create new ones (Price and Boutin 1993), suggesting that the population density is a key ecological agent of selection (Dantzer et al. 2013; Taylor et al. 2014). In most cases, juveniles leave their natal territory in search of vacant territories, ranging on average around 90m, although occasionally up to 900-1000m away from the natal territory (Price and Boutin 1993; Larsen and Boutin 1994; Berteaux and Boutin 2000). However, in some cases the mother will “bequeath” all or part of her territory to one of her offspring, typically a daughter, and search for a vacant territory herself (Price and Boutin 1993; Larsen and Boutin 1994; Berteaux and Boutin 2000; Lane et al. 2015).

Mean litter size in red squirrels is between three and four but can range from one to seven (McAdam et al. 2007). Therefore, there is potential for competition within a litter for maternal resources, nearby available territories, or for access to the mother’s territory if she leaves it. Furthermore, each litter is in competition with the other litters in adjacent territories for vacant territories. Given the distance squirrels can range in search of vacant territories (see above) there is possibly selection at greater spatial scales, for example amongst the young-of-the-year for the few unoccupied territories in the area covered by several territories (“neighbourhoods”), and for competition among neighbourhoods for access to vacant territories within a study area (a rectangular grid of around 40 hectares, here representing a sub-population). Finally, within each year the population is comprised of multiple study areas, so there is possibly selection among these large spatial scales. This creates the opportunity to investigate the strength of selection at different spatial scales: within-litters, within-social neighbourhoods, within-study areas and within-years (amongst-study areas in each year). As claiming a vacant territory is our suggested mode of competition (Taylor et al. 2014), we investigated selection on two traits that are relevant to this ability: birth date and growth rate. Earlier born litters presumably are able to start searching for vacant territories earlier than later ones (Réale et al. 2003a; Williams et al. 2014). A fast growth rate might mean individuals of a given age have an advantage in terms of size when competing for a vacancy (McAdam and Boutin 2003b).

We pursued three main questions. First, what is the strength of selection on growth rate and birth date at each of these levels? Ranking each of these levels of selection also allowed us to identify which was most important to red squirrels. We hypothesized that since settlement distance is typically short (see above), selection will be strongest at the most local scales (i.e. within-litters and within-social neighbourhoods). We also compared this multilevel approach to a standard selection analysis, where we regressed recruitment on individual growth rates and birth dates relative to the yearly average. Secondly, we sought to determine whether, and at what scale, a putative agent of selection, the population density of the study area, affected the direction and magnitude of natural selection. We hypothesized that selection would be intensified by increased population density, although we did not predict which scale would show the most density-dependent selection. Third, as sex-biased patterns of bequeathal may influence selection strengths, we investigated whether these levels of selection differed between males and females. We did not have any previous expectations for which sex would experience stronger selection.

## Materials and Methods

### Study system

We collected data on a wild population of red squirrels in the southwest Yukon, Canada (61° N, 138° W). We have monitored two adjacent study sites (ca. 40 hectares each), bisected by the Alaska highway, continuously since 1987. For this study, we restricted our analyses squirrels born from 1989-2015, as 2015 was the last cohort for which survival data were available. Each year, we live-trapped new individuals (Tomahawk Live Trap, Tomahawk, WI, USA) and gave them unique ear-tags, identified females with litters and ear-tagged their pups, and conducted censuses (using complete enumeration) to ascertain the location and survival of individuals. See McAdam et al. (2007) for further details. These study sites are patches of good habitat among poorer habitat, and hence are somewhere between “islands” and arbitrary areas within a continuous range. As red squirrels can live in the surrounding area, we do see a very low degree of successful emigration from the study area. However, estimated juvenile survival does not differ between the core and the periphery of the study areas, indicating rates of dispersal outside of the study areas are not biasing mortality estimates (McAdam et al. 2007).

Female red squirrels typically give birth to litters between March and May. Young are weaned at approximately 70 days of age (Larsen and Boutin 1994), after which the pups disperse in search of vacant territories or the mother may bequeath a portion or all of her territory to one of her pups (Price and Boutin 1993; Larsen and Boutin 1994; Berteaux and Boutin 2000).

### Data collection

To start monitoring pups as soon as they were born, we regularly live-trapped all females and examined their abdomens and nipples for signs of swelling. We estimated birth date for each litter based on female stages of pregnancy during live-capture events and the size of pups once we found them. For each mother we only used the first litter of the year to allow better comparison among years, as second and third litters are typically only attempted in “mast” years, in which white spruce (*P. glauca*) produces orders of magnitude more seed (Kelly 1994; Boutin et al. 2006; Lamontagne and Boutin 2007) or after failed first litter attempts (McAdam et al. 2007; Williams et al. 2014). To determine their growth rate, we weighed pups twice while they were still within their natal nest, once at 1-2 days old and again at about 25 days old. In this time period their growth is approximately linear (McAdam and Boutin 2003a), so we calculated individual growth rate as the weight difference between the two measures divided by the number of days between the measures, to give growth rate in grams of mass gained per day. We excluded records where the first mass was above 50g, or where the second mass was above 100g, as these were likely to be litters we found late when pup growth rate is no longer linear. We also excluded records when there were fewer than five days between weight measurements. Due to their conspicuous territorial behaviour and our semi-annual censuses of all squirrels, we have nearly perfect knowledge of which squirrels are still alive in the study areas. Each offspring born in the study areas was classified as “recruited” or “did not recruit” based on whether they survived beyond 200 days of age (i.e., survived their first winter). This binary variable was used as the response variable in all our models.

### Data analysis

All analyses were conducted in R ver. 3.3.2 (R Development Core Team 2016), with the package “MCMCglmm” ver. 2.23 (Hadfield 2010). Figures were drawn using coefplot2 (Bolker 2012) and ggplot2 (Wickham 2009).To determine which levels of selection were strongest, we constructed a logistic regression model, containing terms each representing a different level of selection. Therefore, all terms (five for growth rate, four for birth date, see below) were in the same model. The response for the model was the binary variable of whether the individual recruited or not, and we used a logit link function. This meant we were restricted to using absolute rather than relative fitness, but we were still able to calculate selection coefficients, see below. We then calculated each of growth rate and birth date at a series of levels. The first of these for growth rate was the individual’s growth rate relative to the mean of its littermates. This represents within-litter selection. There is no such selection for birth date as all littermates possess the same birth date. The mean of a litter of one was simply the value for the single individual. The next level for growth rate was the mean growth rate of its litter relative to the mean growth rate of all individuals born in nests within 130m of focal nest, representing within-social neighbourhood selection. For birth date we used the birth date of the litter relative to the mean birth date of all litters within its social neighbourhood. The radius of the social neighbourhood was set at 130m, as this is the distance within which squirrels respond to each other’s territorial calls (Smith 1968, 1978), so represents the acoustic social environment an individual experiences. Furthermore, 130m is similar to the distance Dantzer et al. (2012) identified (150m) in this system as being the most relevant for “local” density effects. We repeated the analyses with the social neighbourhood set at 60 or 200m, and found no qualitative differences in the results (see the online supporting information). The next level of selection is within-study area. For this we used the mean growth rate and mean birth date of an individual’s social neighbourhood relative to the mean for the whole study area. We then modelled within-year selection as the mean growth rate and birth date for an individual’s study area relative to the mean growth rate and birth date for the entire year. We also included terms for the year’s mean growth rate and birth date relative to the global mean (across all years and study areas), to control for trait-fitness covariances among-years (e.g. Bouwhuis et al. 2015). Only linear terms were fitted to keep models from getting overly complex and because quadratic terms have previously been shown to be less important than directional selection for these traits in this species (McAdam and Boutin 2003b). This method models an individual’s trait as a series of deviations. For example, an individual with a growth rate of 1.6 g/day might have grown 0.2 g/day slower than the average pup in its litter. This average growth rate of the litter (1.8 g/day) might be 0.3 g/day faster than the average of all litters within the social neighbourhood (1.5 g/day). This may be 0.15g/day slower than the study area-wide mean (1.65g/day) and 0.2g/day slower than the year-wide mean (1.85g/day). This might be 0.1g/day faster than the global mean of 1.75g/day. Therefore, we modelled an individual’s growth rate as the sum of a series of components (1.6 = 1.75 + 0.1 - 0.2 - 0.15 + 0.3 - 0.2), and estimate selection on each using separate partial regression coefficients:

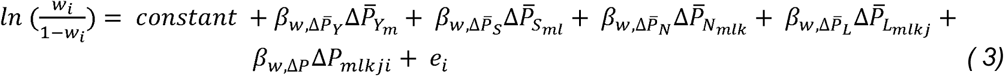

Note as this is a logistic regression we have shown the response variable as the log odds of fitness. 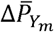 represents the difference between the mean growth rate for the year *m* that *i* was born and the global mean growth rate. 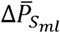 is the difference between the mean growth rate of i’s study area l in year m and the yearly mean. 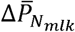 is the difference between the mean growth rate of i’s social neighbourhood k in study area l in year m and the study area mean. 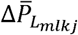 is the difference between the mean growth rate of *i*’s litter *j* in social neighbourhood k in study area l in year m and the neighbourhood mean, and Δ*P*_*mlkji*_ is the difference between i’s growth rate and the mean of its litter j in social neighbourhood k in study area l in year *m*. *βW*… terms are the partial regression coefficients for each component of growth rate on fitness. These logistic regression coefficients were converted into selection coefficients, following Janzen and Stern (1998), to allow comparison with other studies (e.g. Kingsolver *et al.* 2001). This is similar to Bouwhuis et al.’s (2015) analysis on brood mass and survival in great tits (*Parus major*), although for growth rate we have two additional levels (within-social neighbourhood and within-study area). The same formulation was used for birth date, except that there was no within-litter selection. We mean-centred each continuous fixed effect and transformed it by dividing by the variable’s standard deviation, giving each variable a variance of 1. This allowed the effect sizes to be directly compared (Schielzeth 2010). Therefore, by directly comparing the magnitude of the coefficients for each level of growth rate and birth date, we were able to identify the levels at which selection acted most strongly.

Each model also included study area as a fixed effect to control for any variation in survival between the two study areas. We also entered the random effect of year, and the random effects of litter ID nested within mother ID. These accounted for variation in recruitment among years, among litters and among mothers beyond the levels of growth rate and birth date we are studying. As each social neighbourhood was uniquely calculated there was no replication of each social neighbourhood, and so we did not include a random effect for this level. The priors for the variance components followed an inverse-gamma distribution (V = 1, nu = 0.002), with the residual variance fixed at 1, because in a model with a binary response the residual variance is defined by the mean. Models were run for 200,000 iterations, with the first 50,000 discarded and then 1/10 of the remaining iterations used for parameter estimation, to reduce the influence of autocorrelation between successive iterations. Trace plots of the model parameters were checked and a Gelman test for stationarity was used to confirm stable convergence had been achieved (p > 0.156 in all cases). We report the posterior distribution mode (PDM) for each parameter, and the 95% credible intervals (CIs) for this estimate. Our model for the standard selection analysis included individual traits relative to the yearly mean, and the yearly mean relative to the overall mean, as levels of growth rate and birth date. Otherwise the model structure was the same.

### Population density an agent of selection

To test whether population density acted as an agent of selection (Dantzer et al. 2013; Taylor et al. 2014), we took the multilevel model built previously, and added study area population density (number of live adult squirrels per hectare in that study area in that year) as a fixed effect. We interacted this effect with each level of growth rate and birth date in the model, to see how the influence of these competitive traits varied as density changed (Bouwhuis et al. 2015). As before, we mean centred study area density and divided it by the variable’s overall standard deviation. Marginal *R*^2^s (the proportion of total variance explained by the fixed effects) were calculated for each model (Nakagawa and Schielzeth 2013) to determine the change in explanatory power adding our agent of selection had brought.

### Sex-specific selection

We added sex as a fixed effect and the interaction between sex and each level of growth rate and birth date to the first model for multilevel selection (without study area density) to test for sex-specific selection. As sex is a two-level factor, we modelled females as the default and males as a contrast, giving the regression estimate for females and the deviation at each level for males. Note the values for each level of the traits are still relative to the mean of all individuals in the level above, including both sexes.

## Results

Across both study areas in all years (1989-2015) there were 2647 juveniles born that had a known growth rate and birth date at each level. These came from 935 litters from 547 mother squirrels. 26% of these juveniles survived to 200 days. Social neighbourhoods contained a median of four litters (range: 1 – 22) and a median of 11 juveniles (range 1 – 60).

### Levels of selection

Selection on growth rate was positive at all levels, but was strongest within-neighbourhoods and became weaker at both smaller (within-litter selection) and larger hierarchical scales (Fig. 1). There was also a positive among-year effect, such that years with higher growth rate had higher average recruitment. None of the levels of birth date experienced consistent selection, but there was a strong, positive among-year relationship; years where the mean birth date was later had higher recruitment. The was considerable variation among-years in recruitment (PDM = 0.749, CIs = 0.376 to 1.60), essentially no variation among-mothers in recruitment (PDM = 0.02, CIs = <0.001 to 0.350), and a large amount of variation among-litters (PDM = 1.26, CIs = 0.744 to 1.98). There was no difference in juvenile recruitment between the two study areas (PDM = -0.164, CIs = -0.471 to 0.194). The standard selection analysis indicted positive selection on growth rate (PDM = 0.330, CIs = 0.130 to 1.25) but no overall selection on birth date (PDM = -0.066, CIs = -0.198 to 0.089). From Fig. 1 it is apparent that these values represent an aggregation of the different levels of the multilevel analysis.

**Figure 1.**
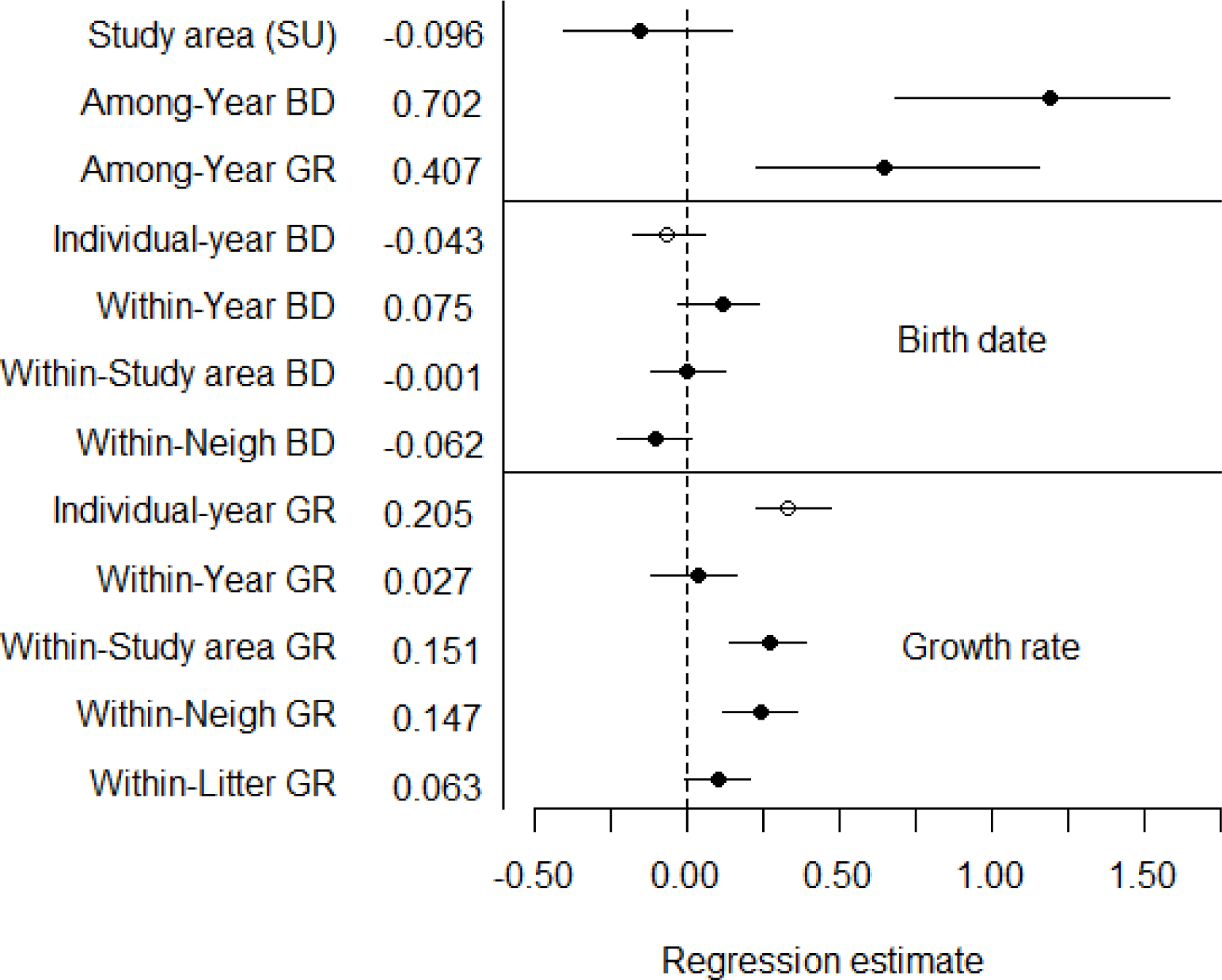
Regression estimates and their 95% credible intervals for the influence of different levels of growth rate (GR) and birth date (BD) on the recruitment of juvenile red squirrels. Also given are the selection coefficients for each trait, obtained following Janzen and Stern (1998). Estimates from the multilevel analysis are indicated with solid points, while the estimates from the standard selection analysis (“Individual-year” terms) are indicated with open circles. Continuous variables have been transformed to the same scale, so effect sizes and selection coefficients are directly comparable. Study area is modelled as a two-level factor, with “Kloo” as the default, and so the effect here shows the difference in the “Sulphur” (SU) study area.

### Agent of selection

Years with high population density experienced stronger within-neighbourhood selection for earlier birth dates. To a lesser degree, within-study area selection on birth date also increased with population density. Within-year selection on birth date, and all levels of selection on growth rate did not vary with changing population density (Table 1). For the majority of our traits (7/9), increasing density increased the strength of selection, as the coefficient for the interaction was of the same sign as for the main effect. However, only for within-neighbourhood selection on birth date did the interaction term not overlap with zero, although the interaction for within-study area selection on birth date only marginally overlapped zero. Adding the fixed effect of study area density, and its interaction with all levels of growth rate and birth date, improved the model fit by 42% (without study area density model R^2^ = 0.144, with study area density model R^2^ = 0.204).

**Table 1.** Posterior distribution mode (PDM) for the estimate of the main effect of each level of growth rate and birth date, and the PDM for the interaction with each effect and study area adult squirrel density (with 95% credible intervals [CIs] in parentheses). Effects for which the CIs did not cross zero are highlighted in bold. When the trait main effect and the interaction between density and the trait act in the same direction then increased density resulted in stronger selection.

| Trait | Effect | PDM of main effect | PDM of interaction | Same direction? |
| --- | --- | --- | --- | --- |
| **Growth rate** | Within-litters | 0.094 (-0.029 to 0.226) | -0.114 (-0.260 to 0.066) | No |
|  | Within-neighbourhoods | **0.232** (0.094 to 0.383) | 0.022 (-0.159 to 0.169) | Yes |
|  | Within-study areas | **0.239** (0.030 to 0.425) | 0.007 (-0.194 to 0.223) | Yes |
|  | Within-years | 0.021 (-0.169 to 0.228) | 0.103 (-0.181 to 0.384) | Yes |
|  | Among-years | **0.694** (0.156 to 1.20) | -0.287 (-0.806 to 0.294) | No |
| **Birth date** | Within-neighbourhoods | **-0.174** (-0.359 to -0.029) | **-0.214** (-0.476 to -0.002) | Yes |
|  | Within-study areas | -0.131 (-0.288 to 0.095) | -0.184 (-0.407 to 0.051) | Yes |
|  | Within-years | 0.169 (-0.057 to 0.340) | 0.104 (-0.205 to 0.331) | Yes |
|  | Among-years | **1.15** (0.534 to 1.71) | 0.091 (-0.402 to 0.596) | Yes |

### Sex-specific selection

Females were more likely to recruit than males (PDM = -0.747, CIs = -1.04 to -0.480; Figs. 2-4). Females that grew faster than their littermates were more likely to recruit, while males were under very little selection for growth rate at this level (Fig. 2a; PDM = -0.403, CIs = - 0.740 to -0.163). Males and females were under equivalent selection for growth rate within-social neighbourhoods (Fig. 2b; PDM = -0.023, CIs = -0.314 to 0.211), within-study areas (Fig. 2c; PDM = -0.117, CIs = -0.415 to 0.107), and within-years (Fig. 2d; PDM = -0.032, CIs = -0.356 to 0.240). The among-year relationship between mean year growth rate and recruitment was positive in females, but tended to be weaker in males (Fig. 3a; PDM = - 0.407, CIs = -0.656 to 0.064). Males and females were under equivalent selection within-social neighbourhoods for birth date (Fig. 4a; PDM = 0.053, CIs = -0.186 to 0.326). Females from neighbourhoods with earlier mean birth dates tended to be more likely to recruit, but the reverse was true for males (Fig. 4b; PDM = 0.311, CIs = 0.021 to 0.528). Males and females were under equivalent selection for birth date within-years (Fig. 4c; PDM = 0.024, CIs = -0.284 to 0.272), but females showed a marginally stronger association between growth rate and recruitment among-years (Fig. 3b; PDM = -0.297, CIs = -0.657 to 0.061). Sex-specific regression estimates are plotted in Fig. 5 to aid interpretation.

**Figure 2.**
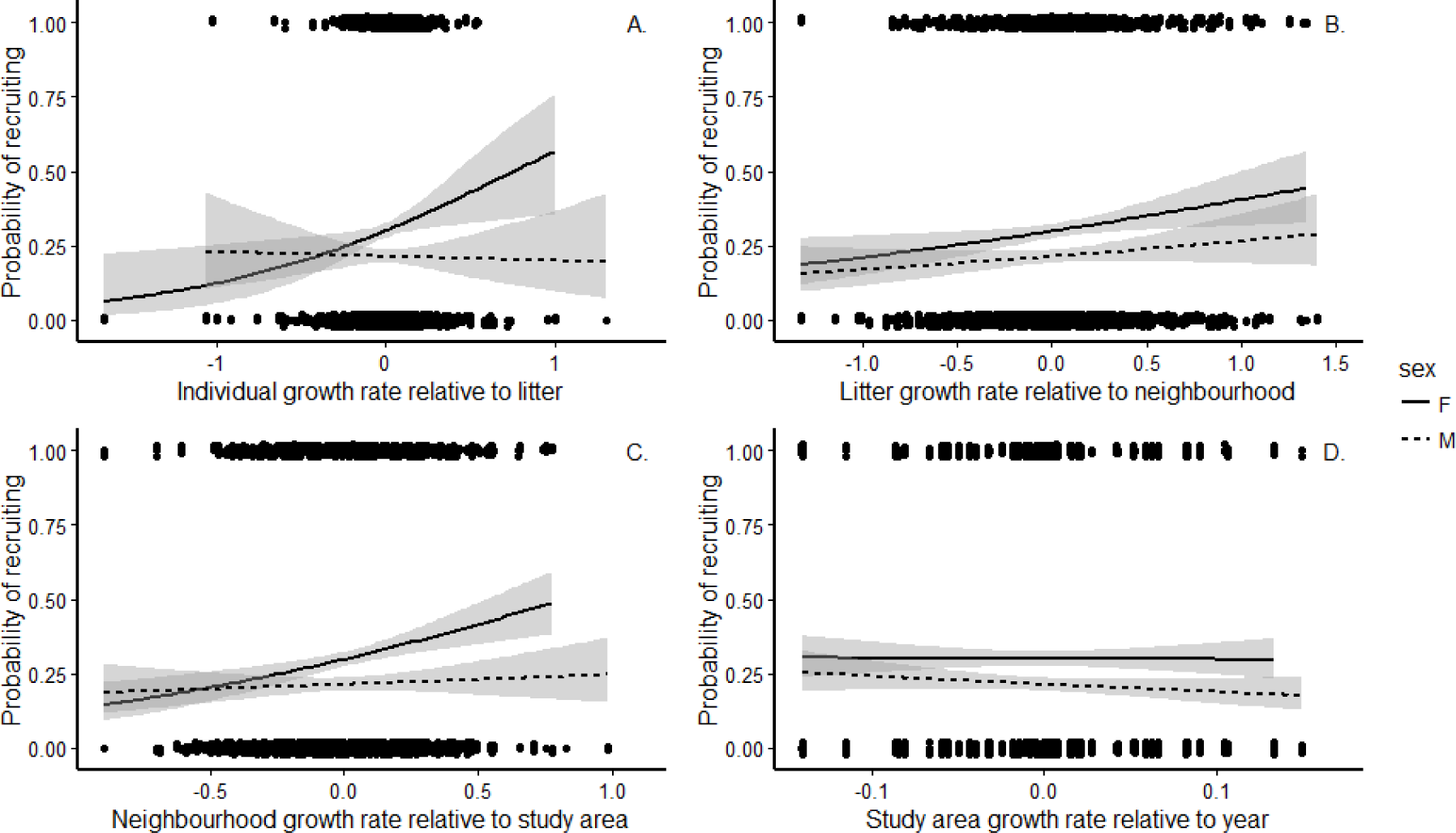
The influence of different levels of growth rate on juvenile red squirrel recruitment. A: Individual growth rate relative to the litter’s mean growth rate. B: Litter mean growth rate relative to the social neighbourhood’s mean growth rate. C: Mean social neighbourhood growth rate relative to the study area’s mean growth rate. D: Study area mean growth rate relative to the mean growth rate for that year. Predictions from the model for females are plotted as a solid line, for males as a dashed line, with the grey areas indicating the standard errors around the estimates. Points have been moved a small amount at random either up or down the y-axis to aid viewing, but all were either 0 or 1.

**Figure 3.**
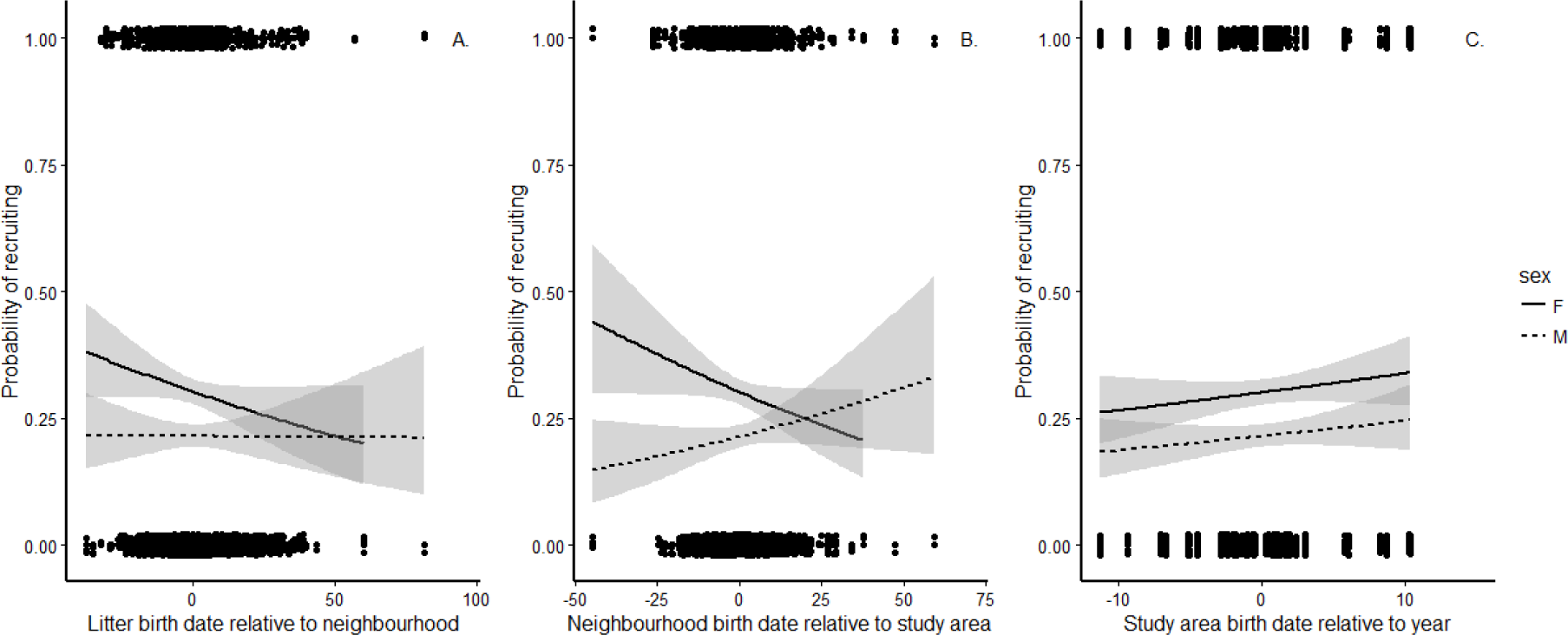
Among-year effects of A: growth rate, and B: birth date, on juvenile red squirrel survival. Predictions from the model for females are plotted as a solid line, for males as a dashed line, with the grey areas indicating the standard errors around the estimates. Points have been moved a small amount at random either up or down the y-axis to aid viewing, but all were either 0 or 1.

**Figure 4.**
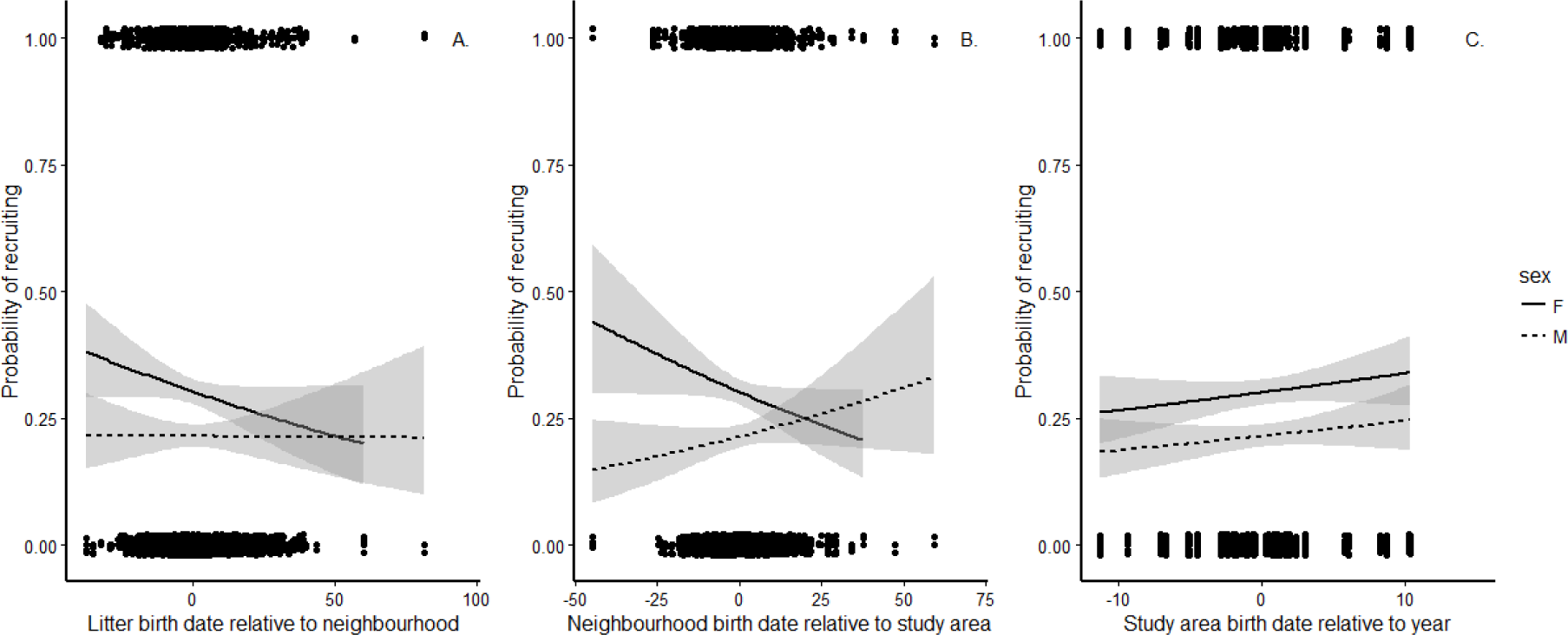
The influence of different levels of birth date on juvenile red squirrel recruitment. A: Litter birth date relative to the social neighbourhood’s mean birth date. B: Mean social neighbourhood birth date relative to the study area’s mean birth date. C: Study area mean birth date relative to the mean birth date for that year. Predictions from the model for females are plotted as a solid line, for males as a dashed line, with the grey areas indicating the standard errors around the estimates. Points have been moved a small amount at random either up or down the y-axis to aid viewing, but all were either 0 or 1.

**Figure 5.**
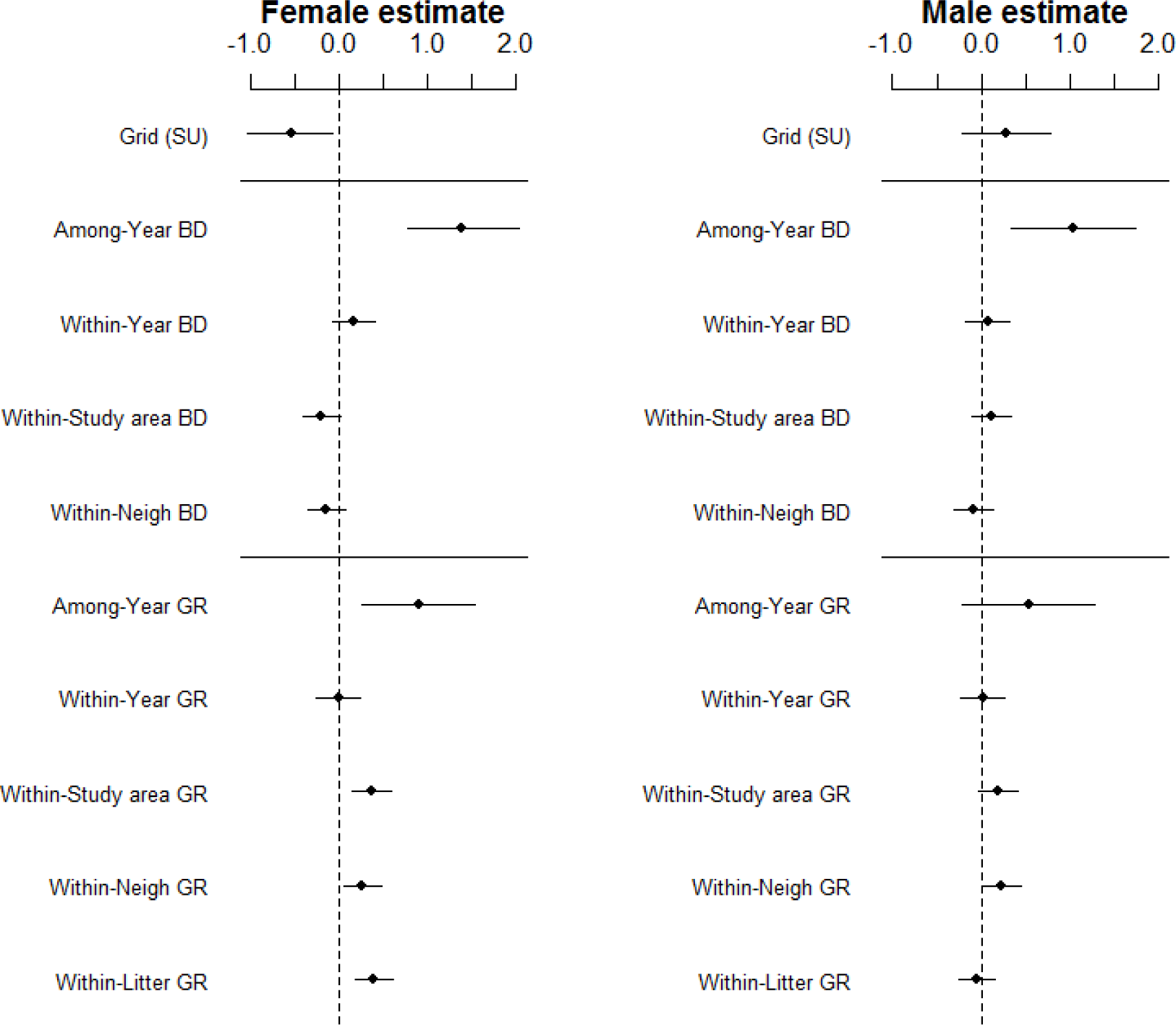
Regression estimates and their 95% credible intervals for the influence of different levels of growth rate (GR) and birth date (BD) on the recruitment of female (left plot) and male (right plot) juvenile red squirrels. Note these were modelled in one model using a sex interaction term, but are plotted here as separate estimates for clarity. Variables have been transformed to the same scale, so effect sizes are directly comparable. Study area is modelled as a two-level factor, with “Kloo” as the default, and so the effect here shows the difference in the “Sulphur” (SU) study area.

## Discussion

### Multilevel-selection

Natural selection on red squirrel growth rates and birth dates was most prominent for both traits within-social neighbourhoods. Being born earlier than neighbouring litters, and/or growing faster than them increased the chances of juveniles recruiting. This level of selection is above the level of the individual squirrel yet is much more local than selection acting across the entire population. Pups who grew faster than their littermates, and from social neighbourhoods that grew faster than others in the study area, were also more likely to recruit. Consistent selection on birth date was only apparent when we added our putative agent of selection, study area density, to the model, indicating that an earlier birth date is primarily beneficial for recruitment when there are many other competing individuals. Therefore, there are interactions among-litters, within a social neighbourhood that are important for whether a juvenile red squirrel recruits or not, and these interactions increase in importance when population density is higher. Consistent selection on birth date was also not apparent from our standard selection analysis, as this value represents an aggregation of the within-and among-study area effects, which were in opposite directions. In contrast, the standard selection analysis did reveal consistent selection favouring faster growth. Our multilevel selection approach revealed that this overall selection was primarily driven by selection acting at the more local scales.

That the within-neighbourhood scale was the most important (although for females within-litter selection on growth rate was stronger, see below) suggests differences among-litters within the social neighbourhood has the largest influence on recruitment in juvenile red squirrels. An evolutionary response to group selection such as this requires non-zero relatedness among-group members (*r* > 0), or alternatively for there to be IGEs among individuals (Bijma and Wade 2008). Litters have a non-zero *r* (mean of between 0.25 and 0.5 depending on the number of fathers, notwithstanding any inbreeding) and as such selection among litters can be expected to result in an evolutionary response. Indeed, previous research has indicated that the majority of evolutionary potential in our system appears to be through selection on litter-level characteristics and indirect maternal effects on these characteristics, as this is where the genetic variance in fitness is (McFarlane et al. 2015) and where selection is strongest (this study, see also: McAdam et al. 2002; McAdam and Boutin 2004). We also note that the response to selection will be influenced by these maternal effects and their correlations with other components of maternal fitness (Thomson et al. 2017), which we have not estimated here. Future studies and predictions on the evolutionary potential of this population should take this in account, as models of evolutionary change incorporating such indirect effects can lead to counter-intuitive results (Mousseau and Fox 1998; Wolf et al. 1998; Bijma and Wade 2008).

Within social-neighbourhood selection being more important than within-study area selection suggests that our definition of a social neighbourhood as all individuals within 130m reflects the level at which red squirrels compete for space and resources to recruit. Further, this is congruent with the work of Dantzer et al. (2012), who demonstrated that density within 150m was the most relevant measure in this system. Red squirrels can hear territorial vocalisations by others from up to 130m (Smith 1968), and mothers use these vocalisations to assess local density and increase the growth rate of their pups through stress-mediated maternal effects (Dantzer et al. 2013). The within-neighbourhood scale did not correspond to a discrete and mutually exclusive ‘group’, but instead represented the unique interactions between each individual and its surrounding neighbours. We add to the results of Laiolo and Obeso (2012) to show that this form of selection can occur based on individually unique social environments, rather than discrete units such as a unique pair or colony (see also: Nunney 1985). For all territorial animals, and those that live in hierarchically structured populations, groups of competing or cooperating animals exist at different scales (Hill et al. 2008). These can be relatively clearly defined, such as a population containing distinct clans formed by discrete family units as found in sperm whales (*Physeter macrocephalus*; Cantor et al. 2015), or defined based on spatial scale as we have done in the current study. Therefore, multilevel selection may be widespread in situations where it has yet to be considered. Genetic relatedness within a social neighbourhood or IGEs among neighbours is required for among-neighbourhood selection to produce a response (Bijma and Wade 2008). Juvenile red squirrels typically do not disperse far from the natal nest (mean around 90m; Price and Boutin 1993; Larsen and Boutin 1994; Berteaux and Boutin 2000), which could lead to clusters of related individuals. Explicit calculation of this parameter will allow us to predict the response to this level of selection.

### Study area density as an agent of selection on birth date

Our putative agent of selection, the density of the study area, was important in determining the strength of selection on birth date at the within-social neighbourhood level, and to a lesser extent the within-study area level, although not for growth rate at any level. Being born earlier than neighbouring litters increased survival, which was especially important when the study area was at a high density, but was less important when density was low. This strengthens the idea that an early birth date is selectively advantageous because it allows juveniles to locate vacant territories within their social neighbourhood.

While previous studies have shown that local density is often negatively related to fitness components (e.g. Coulson et al. 1997; Wilkin et al. 2006), we have identified a trait whose effects on fitness are mediated by population density (MacColl 2011; see also: Dantzer et al. 2013; Bouwhuis et al. 2015). Although our initial analysis suggested no consistent selection on birth date, adding population density to the model revealed both that early-born litters were more likely to recruit, and that this effect was stronger at higher densities. This is likely because there is among-year variation in the strength of selection, related to changes in population size (McAdam and Boutin 2003b), so by accounting for this we were able to detect the effect. Birth date is moderately heritable (h^2^ = 0.16; Réale et al. 2003) and so as predicted by the breeder’s equation should be advancing (Lande 1979). However, despite initial results suggesting a genetic change occurred over a 10-year period (Réale et al. 2003b) additional data and a re-analysis indicated no change in birth date (Lane et al. *In rev*), which seems to be caused by selection acting on environmental deviations rather than the genetic basis to birth date.

### Selection on growth rate

In our analysis, population density was not an agent of selection on growth rate. Dantzer et al. (2013) previously found that a female’s reproductive success was increased if her litter was fast growing when local density was high, but not when it was low, in contrast to our results. They used relative fitness rather than raw survival as their response variable, which shows higher variance when recruitment is lower, which occurs in high-density years. This may have enabled them to detect stronger selection on growth rate in high density years where we did not. In addition, Dantzer et al. (2013) also included litter size in their selection analysis, whereas we included only growth rate and birth date. The degree of competition for vacant territories depends on both the number of vacancies as well as the number of potential competitors (Taylor et al. 2014). While population density represents the inverse of territory vacancy rates, the number of juveniles competing for each vacant territory might also depend on the availability of food resources affecting the rate of offspring production. This mechanism remains to be tested.

Goodnight et al. (1992) stated that if both individual and group-level selection coefficients are the same, the selection is “hard”. The absolute value of the individual’s trait is selected upon, unrelated to the social environment, with the agent likely to be some environmental factor (Goodnight et al. 1992). Considering the selection coefficients were all the same direction for growth rate, and that population density did not greatly influence the strength of selection, selection on growth rate may act in this way. Possibly, faster growing pups are generally of higher “quality”, and so more likely to survive over winter. This too is a mechanism that remains to be tested. Note that the overlapping CIs for the selection coefficients is not necessarily good evidence that selection at different scales is equivalent, as selection strengths fluctuate across years (McAdam and Boutin 2003b).

Although our standard selection analysis indicated strong selection on growth rate, some of this selection occurred at the within-study area level. Response to this section requires genetic variance within-years (among-study areas), which we do not believe is likely. Therefore, this portion of the selection gradient will not contribute to any evolutionary response. This may be a common phenomenon, where standard selection analyses assume that all the selection measured is aligned with the available genetic variation. Our results suggest that might not be the case, which may contribute to the lack of evolutionary response observed in populations where directional selection has been estimated on a heritable trait (Merilä et al. 2001). A thorough multilevel quantitative genetic analysis would be required, however, to completely determine how the scale of selection and the scale of genetic variation together affect rates of evolution of growth rates and birth dates.

### Sex-specific selection at the level of the litter for growth rate

Combining multilevel and sex-specific selection revealed contrasting relationships within-litters for selection on growth rate. Females were under strong, positive selection within the litter, while males were under no selection at this level. Furthermore, females typically were more likely to recruit than males, a relatively common pattern in birds and mammals (Clutton-Brock et al. 1985), and one that has been detected previously in this system (LaMontagne et al. 2013). We suspect that selection was strong within-litters for females as red squirrel mothers sometimes (19% of mothers; Lane et al. 2015) bequeath their territory, or part of it, to one of their offspring (Price and Boutin 1993; Larsen and Boutin 1994), and this offspring is most commonly a daughter (Berteaux and Boutin 2000). If squirrels do disperse from the natal territory, the distance of settlement is not typically very large (see above), and does not differ between the sexes (Cooper et al. *In rev*). Therefore, growing more quickly than its littermates to obtain a larger size is perhaps important for a female squirrel to out-compete its littermates for either the natal territory, or one of the (likely few) available territories near to the nest. As bequeathal is biased towards females, fast growing males may have no better chance of acquiring the natal territory than slower growing males, as the territory tends to go to a female regardless. This may explain the lack of selection for growth rate in males within-litters. Berteaux and Boutin (2000) found that individuals having a territory bequeathed to them were not heavier than those that did not, however this was a population-level analysis, with a smaller sample size than ours, and so may have failed to identify this level of within-litter competition. Alternatively, fast-growing females may have been smaller at birth, but grew more quickly than their siblings. This, however, would oppose the general pattern that individuals that experience catch-up growth suffer reduced longevity (Lee et al. 2012). Young and Badyaev (2004) noted that sex-biased allocation of parental resources is more common when parents are limited in their ability to acquire or store resources. While red squirrels do not appear limited in their ability to store resources, in most years they will be strongly limited in their ability to acquire resources. In mast years this is unlikely to be true. Sex-biased allocation of resources depends on changes in the cost differential of sons and daughters across different environments (Young and Badyaev 2004). Such a cost differential change is not obvious in red squirrels at present, but could be explicitly tested.

We note that the absolute growth rate of individuals did not differ between the sexes (1.73 and 1.75 g/d for females and males respectively; t-test, t = -0.821, df = 2392, p = 0.41), suggesting this selection has not resulted in the evolution of sex-biased growth rates. Sexually antagonistic selection is quite common (Cox and Calsbeek 2009), for instance, some *Anolis* lizard species show sexual eco-morph divergence so that the sexes occupy different ecological niches (Butler et al. 2000, 2007), while body size in female yellow pine chipmunk (*Tamias amoenus*) was typically positively related to fitness, but was selectively neutral in males (Schulte-Hostedde et al. 2002). Sexually antagonistic selection is not necessarily absent in sexually monomorphic species such as the red squirrels, as a sex-specific response may not be possible (Cox and Calsbeek 2009). Although viability selection typically shows the least degree of sexual antagonism (Cox and Calsbeek 2009), we still found evidence for sexually antagonistic selection on recruitment. Similar results have been found in *Drosophila melanogaster*, where when selection on females is prevented, populations evolved towards a slower rate of growth that is favoured in males (Prasad et al. 2007). Cox and Calsbeek (2009) noted that many studies either focus on only one sex, or pool the sexes, despite the fact that sexually antagonistic selection can strongly constrain evolution. Therefore, we can only agree with their assertion that more studies should look for sex-specific patterns of selection. Intriguingly, this sexually antagonistic selection was not apparent at any other level we considered or in previous individual-based selection analyses for these traits (e.g. McAdam and Boutin 2003b). Therefore, considering both sex-specific selection and multilevel selection simultaneously may be necessary in future selection analyses.

### Selection on birth date is opposite at local scales vs. among-years

Offspring from litters born earlier than others in their social neighbourhood had an increased chance of recruitment, yet the among-year effect was in the opposite direction: years that have on average later birth dates had higher mean recruitment. This lead to the standard selection analysis suggesting very limited selection on birth dates. This among-year effect is driven by annual variation in resource abundance. In mast years, litters tend to be born later (Boutin et al. 2006). The recruitment in these years is then increased as there are far more resources available, allowing juveniles to create territories where there were none previously and cache food there, increasing survival over winter (McAdam and Boutin 2003b). We also note that selection on growth rates is stronger in the year *after* one of high cone abundance (i.e. after a mast year), likely due to high densities, but that episodes of strong selection are rare (McAdam and Boutin 2003b). Therefore, consistent within-year selection may not always be apparent if among-year variation is not accounted for. Among-year relationships between environmental conditions and reproductive dates alongside selection within each year for these dates to shift earlier have been found in collared flycatchers (*F. albicollis*) and red deer (*Cervus elaphus*), where females alter reproductive dates based on local temperature or previous autumn rainfall respectively (Brommer et al. 2005; Nussey et al. 2005). Therefore, the masking of within-year selective forces by among-year variance in environmental conditions may be common, and so controlling for it necessary when investigating selection (see also van de Pol and Wright 2009 for analogous within-and among-individual effects).

### Conclusions

We have detected multilevel selection on recruitment in a natural population of red squirrels. Selection was typically strongest when considering all individuals within the acoustic social neighbourhood, although females also experienced strong within-litter selection on growth rate. We also found evidence that population density acted as an agent of selection on birth date during juvenile recruitment, but we found no evidence of density-dependent selection through growth rate. If, as our results suggest, interactions are strongest at the within-neighbourhood level, then evolutionary dynamics will strongly depend on traits and genetic parameters at this level, alongside the individual level (Goodnight et al. 1992; Bijma and Wade 2008). Our results highlight 1) the range of scales at which natural selection might act in a solitary organism, 2) how identifying the agent of selection helps us understand a system, 3) that sex-specific selection can occur only at particular levels of organisation, and 4) coefficients of selection being in the same or opposite direction across levels may lead to the over-or under-estimation of selection. A better understanding of how natural selection acts across a range of scales will enhance our ability to understand and predict trait evolution in natural populations.

## Acknowledgements

The authors are grateful for funding from the Natural Sciences and Engineering Research Council, the Northern Scientific Training Program, the National Science Foundation, and the Ontario Ministry of Research and Innovation. We thank Agnes Moose and her family for long-term access to her trapline, and to the Champagne and Aishihik First Nations for allowing us to conduct work on their land. We thank all the volunteers, field assistants and graduate students whose tireless work makes the KRSP possible. We have no conflicts of interest. This is KRSP paper number 83.

## References

Arnold, S. 1983. Morphology, Performance and Fitness. Am. Zool. 361:p347–361.

Berteaux, D., and S. Boutin. 2000. Breeding dispersal in female North American red squirrels. Ecology 81:p1311–1326.

Biernaskie, J. M., and K. R. Foster. 2016. Ecology and multilevel selection explain aggression in spider colonies. Ecol. Lett. 19:p873–879.

Bijma, P., W. M. Muir, and J. A. M. Van Arendonk. 2007. Multilevel selection 1: Quantitative genetics of inheritance and response to selection. Genetics 175:p277–88.

Bijma, P., and M. J. Wade. 2008. The joint effects of kin, multilevel selection and indirect genetic effects on response to genetic selection. J. Evol. Biol. 21:p1175–88.

Björklund, M., and L. Gustafsson. 2013. The importance of selection at the level of the pair over 25 years in a natural population of birds. Ecol. Evol. 3:p4610–4619.

Bolker, B. 2012. coefplot2.

Boutin, S., L. A. Wauters, A. McAdam, M. Humphries, G. Tosi, and A. Dhondt. 2006. Anticipatory reproduction and population growth in seed predators. Science (80-.). 314:p1928–1930.

Bouwhuis, S., O. Vedder, C. J. Garroway, and B. C. Sheldon. 2015. Ecological causes of multilevel covariance between size and first-year survival in a wild bird population. J. Anim. Ecol. 84:p208–218.

Brommer, J. E., J. Merilä, B. C. Sheldon, and L. Gustafsson. 2005. Natural selection and genetic variation for reproductive reaction norms in a wild bird population. Evolution (N. Y). 59:p1362–1371.

Butler, M. A., S. A. Sawyer, and J. B. Losos. 2007. Sexual dimorphism and adaptive radiation in Anolis lizards. Nature 447:p202–205.

Butler, M. A., T. W. Schoener, and J. B. Losos. 2000. The relationship between sexual size dimorphism and habitat use in Greater Antillean Anolis lizards. Evolution 54:p259–72.

Cantor, M., L. G. Shoemaker, R. B. Cabral, C. O. Flores, M. Varga, and H. Whitehead. 2015. Multilevel animal societies can emerge from cultural transmission. Nat. Commun. 6:8091.

Clutton-Brock, T. H., S. D. Albon, and F. E. Guinness. 1985. Parental investment and sex differences in juvenile mortality in birds and mammals. Nature 313:p131–133.

Coulson, T., S. Albon, F. Guinness, J. Pemberton, and T. Clutton-Brock. 1997. Population substructure, local density, and calf winter survival in red deer (Cervus elaphus). Ecology 78:p852–863. Ecological Society of America.

Cox, R. M., and R. Calsbeek. 2009. Sexually antagonistic selection, sexual dimorphism, and the resolution of intralocus sexual conflict. Am. Nat. 173:p176–187.

Dantzer, B., S. Boutin, M. M. Humphries, and A. G. McAdam. 2012. Behavioral responses of territorial red squirrels to natural and experimental variation in population density. Behav. Ecol. Sociobiol. 66:p865–878.

Dantzer, B., A. E. M. Newman, R. Boonstra, R. Palme, S. Boutin, M. M. Humphries, and A. G. McAdam. 2013. Density Triggers Maternal Hormones That Increase Adaptive Offspring Growth in a Wild Mammal. Science (80-.). 340:p1215–1217.

Eldakar, O. T., M. J. Dlugos, J. W. Pepper, and D. S. Wilson. 2009. Population structure mediates sexual conflict in water striders. Science 326:816.

Eldakar, O., D. Wilson, M. Dlugos, and J. Pepper. 2010. The role of multilevel selection in the evolution of sexual conflict in the water strider Aquarius remigis. Evolution (N. Y).

Endler, J. A. 1986. Natural selection in the wild. Princeton University Press.

Falconer, D. 1981. Introduction to Quantitative Genetics. The Ronald Press Company, New York.

Gardner, A. 2015. The genetical theory of multilevel selection. J. Evol. Biol. 28:p305–19.

Goodnight, C. J. 2015. Multilevel selection theory and evidence: a critique of Gardner, 2015. J. Evol. Biol. 28:p1734–46.

Goodnight, C. J., J. M. Schwartz, and L. Stevens. 1992. Contextual analysis of models of group selection, soft selection, hard selection, and the evolution of altruism. Am. Nat. 140:743–761.

Goodnight, C. J., and L. Stevens. 1997. Experimental Studies of Group Selection: What Do They Tell US About Group Selection in Nature? Am. Nat. 150:S59–S79.

Grafen, A. 1984. Natural selection, kin selection and group selection.

Hadfield, J. D. 2010. MCMC methods for multi-response generalized linear mixed modelsLJ: The MCMCglmm R package. J. Stat. Softw. 33:1–22.

Heisler, I. L., and J. Damuth. 1987. A Method for Analyzing Selection in Hierarchically Structured Populations. Am. Nat. 130:582.

Hill, R. A., R. A. Bentley, and R. I. M. Dunbar. 2008. Network scaling reveals consistent fractal pattern in hierarchical mammalian societies. Biol. Lett. 4:748–51.

Janzen, F. J., and S. Stern, Hal. 1998. Logistic regression for empirical studies of multivariate selection. Evolution (N. Y). 52:1564–1571.

Kelly, D. 1994. The evolutionary ecology of mast seeding. Trends Ecol. Evol. 9:465–470.

Kingsolver, J. G., H. E. Hoekstra, J. M. Hoekstra, D. Berrigan, S. N. Vignieri, C. E. Hill, A. Hoang, P. Gibert, and P. Beerli. 2001. The strength of phenotypic selection in natural populations. Am. Nat. 157:245–261. The University of Chicago Press.

Laiolo, P., and J. R. Obeso. 2012. Multilevel selection and neighbourhood effects from individual to metapopulation in a wild passerine. PLoS One 7:e38526.

Lamontagne, J. M., and S. Boutin. 2007. Local-scale synchrony and variability in mast seed production patterns of Picea glauca. J. Ecol. 95:991–1000.

LaMontagne, J. M., C. T. Williams, J. L. Donald, M. M. Humphries, A. G. McAdam, and S. Boutin. 2013. Linking intraspecific variation in territory size, cone supply, and survival of North American red squirrels. J. Mammal. 94:1048–1058. American Society of Mammalogists.

Lande, R. 1979. Quantitative genetic analysis of multivariate evolution, applied to brainLJ: body size allometry. Evolution (N. Y). 33:402–416.

Lande, R., and S. Arnold. 1983. The measurement of selection on correlated characters.

Lane, J. E., A. G. McAdam, A. Charmantier, M. M. Humphries, D. W. Coltman, Q. Fletcher, J. C. Gorrell, and S. Boutin. 2015. Post-weaning parental care increases fitness but is not heritable in North American red squirrels. J. Evol. Biol. 28:1203–12.

Larsen, K. W., and S. Boutin. 1994. Movements, survival, and settlement of red squirrel (Tamiasciurus hudsonicus) offspring. Ecology 75:214–223.

Lee, W.-S., P. Monaghan, and N. B. Metcalfe. 2012. Experimental demonstration of the growth rate–lifespan trade-off. Proc. R. Soc. London B Biol. Sci. 280.

Lush, J. 1937. Animal Breeding Plans. Iowa State College Press, Ames, Iowa.

MacColl, A. D. C. 2011. The ecological causes of evolution. Trends Ecol. Evol. 26:514–522.

Mcadam, A. G., and S. Boutin. 2004. Maternal effects and the response to selection in red squirrels. Proc. R. Soc. B Biol. Sci. 271:75–79. The Royal Society.

McAdam, A. G., and S. Boutin. 2003a. Effects of food abundance on genetic and maternal variation in the growth rate of juvenile red squirrels. J. Evol. Biol. 16:1249–1256. Blackwell Science Ltd.

McAdam, A. G., and S. Boutin. 2003b. Variation in viability selection among cohorts of juvenile red squirrels (Tamiasciurus hudsonicus). Evolution 57:1689–1697. Blackwell Publishing Ltd.

McAdam, A. G., S. Boutin, D. Réale, and D. Berteaux. 2002. Maternal effects and the potential for evolution in a natural population of animals. Evolution 56:846–851.

McAdam, A. G., S. Boutin, A. K. Sykes, and M. M. Humphries. 2007. Life histories of female red squirrels and their contributions to population growth and lifetime fitness. Ecoscience 14:362.

McFarlane, S. E., J. C. Gorrell, D. W. Coltman, M. M. Humphries, S. Boutin, and A. G. McAdam. 2015. The nature of nurture in a wild mammal’s fitness. Proc. R. Soc. B Biol. Sci. 282:20142422–20142422.

McGlothlin, J. W., A. J. Moore, J. B. Wolf, and E. D. Brodie. 2010. Interacting phenotypes and the evolutionary process. III. Social evolution. Evolution 64:2558–74.

Merilä, J., B. C. Sheldon, and L. E. Kruuk. 2001. Explaining stasis: microevolutionary studies in natural populations. Genetica 112–113:199–222.

Mousseau, T., and C. Fox. 1998. The adaptive significance of maternal effects. Trends Ecol. Evol.

Muir, W. M., P. Bijma, and A. Schinckel. 2013. Multilevel selection with kin and non-kin groups, experimental results with japanese quail (coturnix japonica). Evolution (N. Y). 67:1598–1606.

Nakagawa, S., and H. Schielzeth. 2013. A general and simple method for obtaining R 2 from generalized linear mixed-effects models. Methods Ecol. Evol. 4:133–142.

Nunney, L. 1985. Group selection, altruism, and structured-deme models. Am. Nat. 126:212–230.

Nussey, D. H., T. H. Clutton-Brock, D. A. Elston, S. D. Albon, and L. E. B. Kruuk. 2005. Phenotypic plasticity in a maternal trait in red deer. J. Anim. Ecol. 74:387–396. Blackwell Science Ltd.

Page, R. E., and M. K. Fondrk. 1995. The effects of colony-level selection on the social organization of honey bee (Apis mellifera L.) colonies: colony-level components of pollen hoarding. Behav. Ecol. Sociobiol. 36:135–144. Springer-Verlag.

Prasad, N. G., S. Bedhomme, T. Day, and A. K. Chippindale. 2007. An evolutionary cost of separate genders revealed by male-limited evolution. Am. Nat. 169:29–37.

Price, G. R. 1970. Selection and covariance. Nature Publishing Group.

Price, K., and S. Boutin. 1993. Territorial bequathal by red squirrel mothers. Behav. Ecol. 4:144–155. Oxford University Press.

Queller, D. C. 1992. Quantitative genetics, inclusive fitness, and group selection. Am. Nat. 139:540.

R Development Core Team, R. 2016. R: A language and environment for statistical computing. R Foundation for Statistical Computing, Vienna, Austria.

Réale, D., D. Berteaux, A. G. McAdam, and S. Boutin. 2003a. Lifetime selection on heritable life-history traits in a natural population of red squirrels. Evolution (N. Y). 57:2416–2423.

Réale, D., A. G. McAdam, S. Boutin, and D. Berteaux. 2003b. Genetic and plastic responses of a northern mammal to climate change. Proc. Biol. Sci. 270:591–596.

Robertson, A. 1966. A mathematical model of the culling process in dairy cattle. Anim. Prod. 8:95–108. Cambridge University Press.

Royle, N. J., N. J. Royle, I. R. Hartley, I. R. Hartley, I. P. F. Owens, I. P. F. Owens, G. A. Parker, and G. A. Parker. 1999. Sibling competition and the evolution of growth rates in birds. Proc. R. Soc. Lond. B 266:923–932.

Schielzeth, H. 2010. Simple means to improve the interpretability of regression coefficients. Methods Ecol. Evol. 1:103–113.

Schulte-Hostedde, A. I., J. S. Millar, and H. L. Gibbs. 2002. Female-biased sexual size dimorphism in the yellow-pine chipmunk (Tamias amoenus): sex-specific patterns of annual reproductive success and survival. Evolution 56:2519–29.

Siepielski, A. M., J. D. Dibattista, and S. M. Carlson. 2009. It’s about time: The temporal dynamics of phenotypic selection in the wild. Blackwell Publishing Ltd.

Siepielski, A. M., K. M. Gotanda, M. B. Morrissey, S. E. Diamond, J. D. DiBattista, and S. M. Carlson. 2013. The spatial patterns of directional phenotypic selection. Ecol. Lett. 16:1382–1392.

Smith, B. R., and D. T. Blumstein. 2008. Fitness consequences of personality: a meta-analysis. Behav. Ecol. 19:448–455. Oxford University Press.

Smith, C. C. 1978. Structure and function of the vocalizations of tree squirrels (Tamiasciurus). J. Mammal. 59:793–808. The Oxford University Press.

Smith, C. C. 1968. The adaptive nature of social organization in the genus of three squirrels Tamiasciurus. Ecol. Monogr. 38:31–64. Ecological Society of America.

Stevens, L., C. J. Goodnight, and S. Kalisz. 1995. Multilevel Selection in Natural Populations of Impatiens capensis. Am. Nat. 145:513–526. University of Chicago Press.

Taylor, R. W., S. Boutin, M. M. Humphries, and A. G. McAdam. 2014. Selection on female behaviour fluctuates with offspring environment. J. Evol. Biol. 27:2308–21.

Thomson, C. E., F. Bayer, M. Cassinello, N. Crouch, E. Heap, E. Mittell, and J. D. Hadfield. 2017. Selection on parental performance opposes selection for larger body size in a wild population of blue tits. Evolution (N. Y). 71:716–732.

van de Pol, M., and J. Wright. 2009. A simple method for distinguishing within-versus between-subject effects using mixed models. Anim. Behav. 77:753–758.

van Veelen, M., J. García, M. W. Sabelis, and M. Egas. 2012. Group selection and inclusive fitness are not equivalent; the Price equation vs. models and statistics. J. Theor. Biol. 299:64–80.

Werschkul, D. F., and J. A. Jackson. 1979. Sibling competition and avian growth rates. Ibis (Lond. 1859). 121:97–102. Blackwell Publishing Ltd.

Wickham, H. 2009. ggplot2: elegant graphics for data analysis. Springer New York, New York.

Wilkin, T. A., D. Garant, A. G. Gosler, and B. C. Sheldon. 2006. Density effects on life-history traits in a wild population of the great tit Parus major: Analyses of long-term data with GIS techniques. J. Anim. Ecol. 75:604–615. Blackwell Publishing Ltd.

Williams, C. T., J. E. Lane, M. M. Humphries, A. G. McAdam, and S. Boutin. 2014. Reproductive phenology of a food-hoarding mast-seed consumer: resource-and density-dependent benefits of early breeding in red squirrels. Oecologia 174:777–788. Springer Berlin Heidelberg.

Wolf, J. B., E. D. Brodie iii, J. M. Cheverud, A. J. Moore, and M. J. Wade. 1998. Evolutionary consequences of indirect genetic effects. Trends Ecol. Evol. 13:64–69.

Wolf, J. B., E. D. Brodie iii, and A. J. Moore. 1999. Interacting Phenotypes and the Evolutionary Process. II. Selection Resulting from Social Interactions. Am. Nat. 153:254–266.

Young, R. L., and A. V. Badyaev. 2004. Evolution of sex-biased maternal effects in birds: I. Sex-specific resource allocation among simultaneously growing oocytes. J. Evol. Biol. 17:1355–1366. Blackwell Science Ltd.

